# A thermoregulatory design principle for transitions into hypometabolism

**DOI:** 10.64898/2026.09.25.754297

**Authors:** Hikaru Sugimoto, Genshiro A. Sunagawa, Satoshi Nakagawa, Takeshi Sakurai, Yoshifumi Yamaguchi, Shinya Kuroda

**Author notes:** Corresponding authors: Genshiro A. Sunagawa, Shinya Kuroda.

## Abstract

Mammals entering torpor or hibernation undergo an abrupt transition from normothermia to hypothermia, yet how thermoregulation enables this switch remains poorly understood. Here, we identify dynamical signatures that precede these transitions and a mathematical principle that can generate them. In fasting-induced torpor in mice, body-temperature fluctuations increased before torpor onset, providing an early-warning signal that tracked proximity to the transition better than temperature decline alone. A heat-balance model showed that reducing how strongly the effective heat-loss coefficient depends on body temperature reorganizes thermoregulatory stability, allowing a low-temperature equilibrium to emerge while the normothermic state remains stable. This organization is consistent with a symmetry-broken pitchfork involving a saddle-node. Similar increases in temperature fluctuations preceded hibernation onset in hamsters. These findings link pre-transition temperature dynamics to changes in the underlying thermoregulatory landscape and provide a framework for detecting and understanding transitions from normothermia to hypothermia.

## Introduction

Efficient regulation of energy expenditure is essential for animals enduring prolonged cold and food scarcity. Among the strategies employed under such conditions are hibernation and torpor, which involve significant reductions in metabolic rate and body temperature^1^. Despite extensive studies motivated by basic physiology and potential biomedical applications^1,2^, the regulatory mechanisms controlling the onset, maintenance, and cessation of these states remain unclear.

Previous studies have characterized the thermal patterns accompanying these energy-saving states^3–16^. Systematic declines in body temperature during pre-hibernation have been reported in various hibernating species^3–5^, including Syrian hamster (*Mesocricetus auratus*), Arctic ground squirrel (*Urocitellus parryii*) and Alpine marmot (*Marmota marmota*). The temporal structure of torpor–arousal cycles appears to be governed by at least two endogenous periods: a multiday rhythm and an annual component^6^. A mathematical model using steady-state data suggested that daily torpor in mice (*Mus musculus*) is associated with minimal shifts in the temperature set point^7^. While these studies elucidate important facets of thermoregulatory physiology, such as pre-hibernation cooling, torpor-arousal cycling patterns, and steady-state properties, the field lacks a quantitative framework that captures the full progression from normothermia to torpor and provides measurable indicators of how close the system is to torpor onset.

Here, we develop a mathematical model of the transition from pre-torpor to torpor. We test the hypothesis that the transition to hypometabolic states is organized by a pitchfork-type organization, in which stable states at lower and higher body temperatures become available beyond a critical parameter threshold; torpor corresponds to a transition toward the lower-temperature branch (**Fig. 1**). We further evaluate whether early warning signals of approaching dynamical transitions^8,17,18^, such as increased variance and autocorrelation associated with weakening restoring dynamics, can indicate proximity to the onset of hibernation and torpor. We also ask whether such signals could arise as the thermoregulatory landscape becomes progressively less resistant to temperature excursions before a low-temperature state becomes accessible.

**Figure 1.**
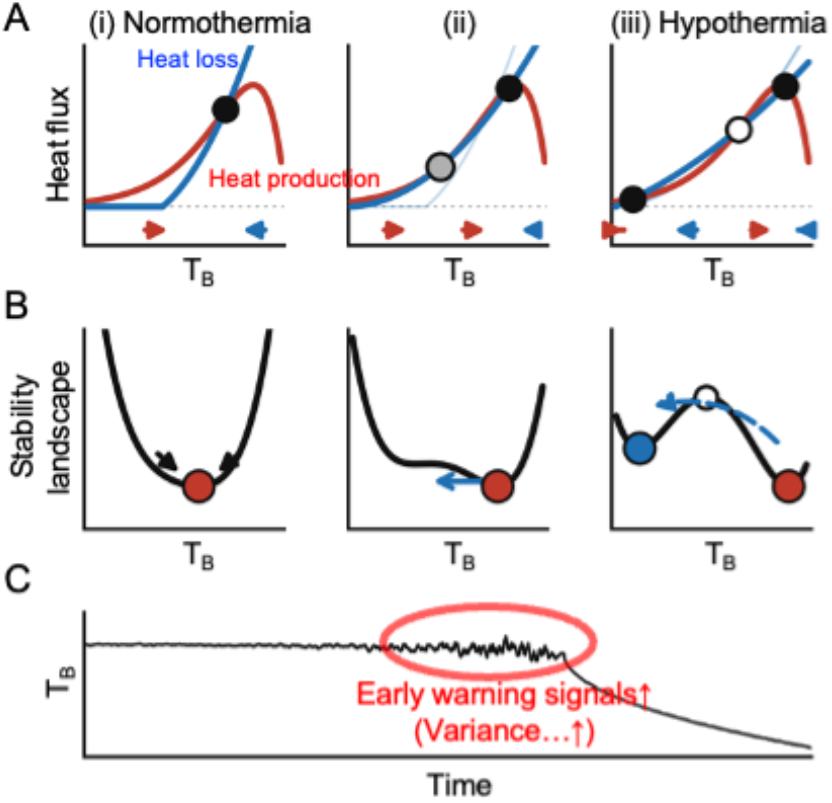
A conceptual schematic of a pitchfork-type organization underlying transitions to torpor/hibernation. (A) Heat production (red) and heat loss (blue) as functions of T_B_. A steady state occurs wherever the two curves cross; filled circles denote stable and open circles unstable steady states, and the arrows beneath each panel give the direction in which T_B_ changes. (i) In the normothermic state the curves cross once, so normothermia is the only steady state. (ii) When the temperature dependence of heat loss becomes shallower, the two curves approach tangency (gray circle); this is the point at which a new pair of steady states is created. (iii) Past that point the curves cross three times, so a second stable state is available at low T_B_. Faint blue curves reproduce the heat-loss curve of the preceding panel. (B) The same three situations drawn as a stability landscape, in which T_B_ behaves like a ball resting in a valley. (i) A deep valley holds T_B_ firmly. (ii) The valley becomes shallow, preferentially on the cold side, so T_B_ fluctuates more and returns to the steady state more slowly; this is what variance and autocorrelation are expected to detect. (iii) A second valley is available at low T_B_, and entry into torpor corresponds to moving into it (dashed arrow). (C) Schematic time course of T_B_: fluctuations grow as the landscape flattens, and T_B_ then falls below the torpor threshold at onset. In the terminology used in the Results, the creation of the new pair of steady states in (A, ii) is a saddle-node bifurcation, and the resulting arrangement of two stable states is that of a symmetry-broken pitchfork; panels A and B are schematics and are not drawn to the scale of the fitted model. Figures 2 and 3 evaluate whether early warning signals (*e*.*g*., increased variance), which typically precede bifurcation points, can be detected before the onset of torpor and hibernation. Figure 4 assesses whether a heat-balance model fitted to mouse data reproduces the change from (A, i) to (A, iii) and can describe the body-temperature (T_B_) dynamics observed in mice.

## Results

### Early warning signals track proximity to fasting-induced torpor in mice

To test whether early warning signal (EWS) metrics change before the onset of fasting-induced torpor, we analyzed core body temperature (T_B_) in adult male C57BL/6J mice (n = 37; 6-8 mice per ambient temperature) across five ambient temperatures (**Fig. 2A, Methods**). All 37 mice entered torpor and were included in the analyses. **Figure 2B** shows representative T_B_ profiles of mice entering torpor. Consistent with the hypothesis, EWS (variance, autocorrelation, and the power-spectrum-based index Smax) showed significant positive Kendall’s τ correlation with time (*Q* < 0.05; **Fig. 2B, C**), indicating progressive increases in EWS prior to torpor. To determine which class of transition the pre-torpor dynamics most resemble, we applied a previously validated deep-learning classifier trained on labeled bifurcation types^17^. Among the candidate classes, the model assigned the highest probability to a pitchfork bifurcation (**Fig. 2D**). The classifier discriminates among several idealized bifurcation normal forms, one of which is a symmetric pitchfork; it therefore cannot by itself establish that the empirical system has an exact pitchfork symmetry or distinguish a pitchfork from a nearby asymmetric unfolding. Moreover, the assigned probabilities were modest in absolute terms and the separation between classes was small. We therefore treated this result as hypothesis-generating rather than as evidence establishing a pitchfork and tested the fitted model structure directly below.

**Figure 2.**
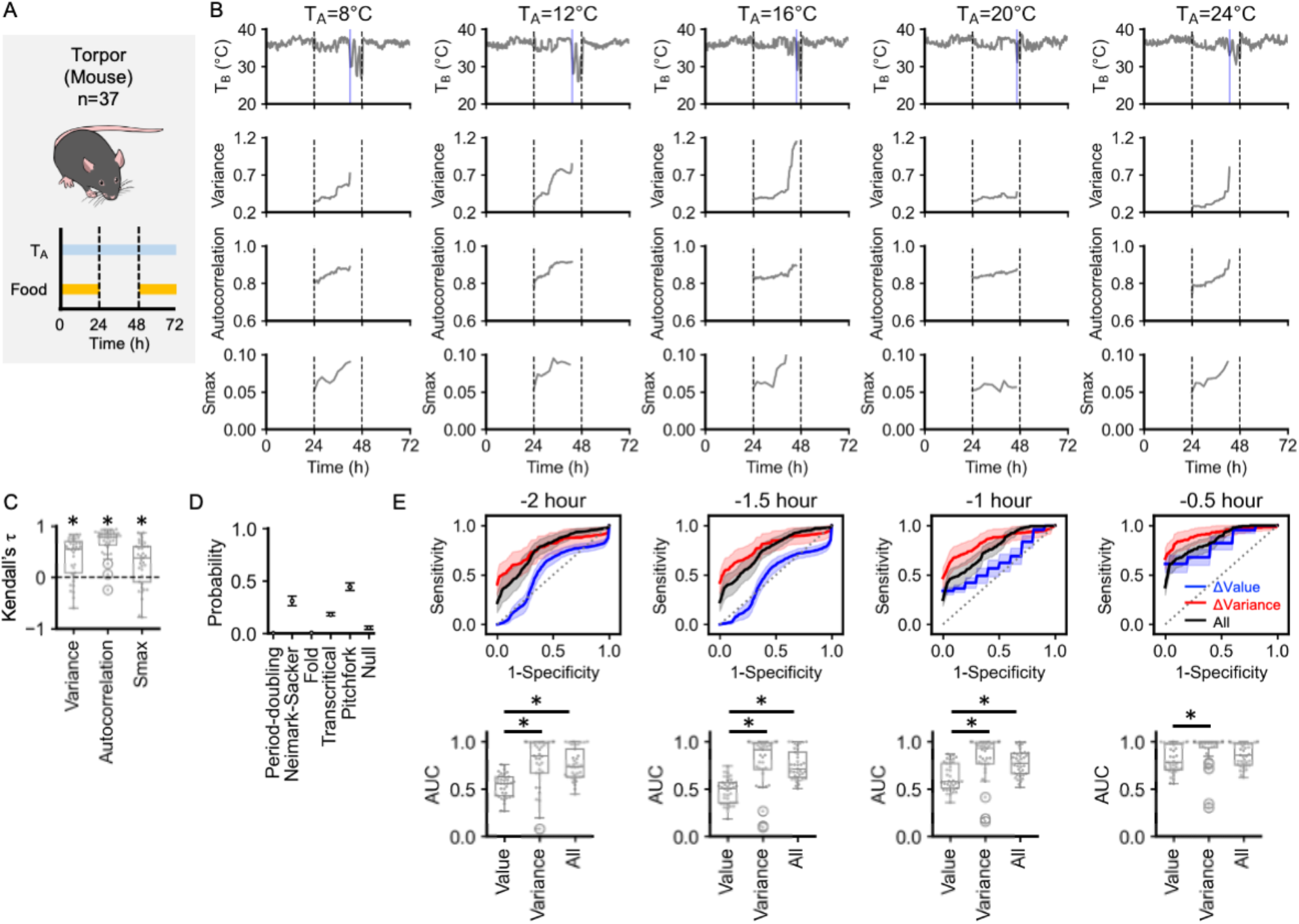
Early-warning signals track proximity to fasting-induced torpor in mice. (A) Experimental design. Mice (n = 37) were housed individually in metabolic chambers under constant ambient temperature (T_A_). Core body temperature (T_B_) was monitored during three 24-h periods: baseline (*ad libitum* food access), fasting (food withdrawn), and recovery (food reintroduction). (B) Representative time series at five T_A_ conditions (8–24 °C): T_B_ (top row) and corresponding early warning signals, including variance, autocorrelation and Smax. Early warning signals were calculated using a 24-hour rolling window until T_B_ dropped below the torpor threshold (33 °C, indicated by blue lines). (C) Box plots showing Kendall’s τ correlation coefficients for each early warning signal. (D) 95% confidence intervals for the probability of different types of bifurcation derived from the machine learning analysis of pre-torpor T_B_ dynamics. (E) Within-trajectory discrimination of proximity to torpor onset at four lead times (2, 1.5, 1, and 0.5 hours). Upper panels: Receiver operating characteristic (ROC) curves. Shaded areas indicate 95% confidence intervals. Lower panels: Box plots of area under the ROC curve (AUC) for each model. Each point corresponds to the value for a single mouse. \**Q* < 0.05.

Next, we assessed the utility of EWS for tracking proximity to torpor onset at lead times of 2, 1.5, 1, and 0.5 hours. A model that used changes in T_B_ variance (ΔVariance) yielded significantly higher within-trajectory areas under the receiver operating characteristic curve (AUCs) than a model that used deviation of T_B_ relative to the first window of the epoch (ΔValue) at every lead time (*Q* < 0.05 for all; **Fig. 2E**). Adding T_B_ deviation to variance (All) did not yield a statistically significant improvement over ΔVariance alone. Collectively, EWS provided information about proximity to torpor onset.

### Early warning signals track proximity to hibernation in Syrian hamsters

We then investigated whether similar signals precede hibernation in Syrian hamsters. The animals (n = 19) were shifted to a short photoperiod and a lower ambient temperature to induce hibernation (**Fig. 3A, B, Methods**). Fourteen of the animals entered hibernation, while five did not. In animals that entered hibernation, variance, autocorrelation, and Smax increased in the days preceding hibernation onset, and the corresponding Kendall’s τ values were significantly higher than in animals that did not enter hibernation (*Q* < 0.05; **Fig. 3B, C**). Using the same classifier as for the mice, we again found that the highest probability was assigned to a pitchfork bifurcation (**Fig. 3D**). As in mice, the assigned probabilities were modest and the confidence intervals of the leading classes overlapped, so this result was likewise treated as hypothesis-generating.

**Figure 3.**
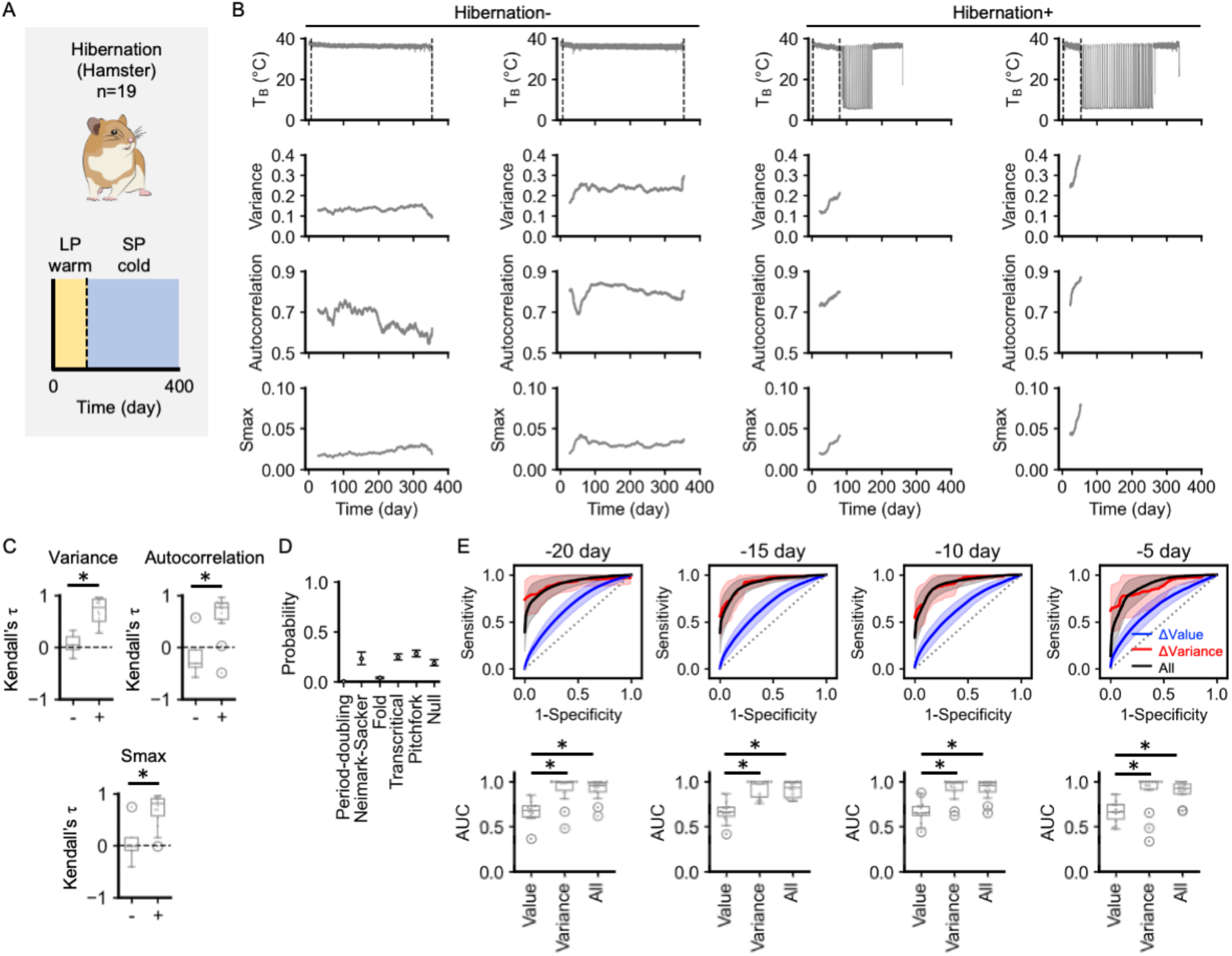
Early-warning signals track proximity to hibernation onset in Syrian hamsters. (A) Experimental design. Syrian hamsters (n = 19) were maintained under summer-like conditions (long photoperiod, LP; warm T_A_) and then transferred to winter-like conditions (short photoperiod, SP; cold T_A_) to induce hibernation. (B) Representative T_B_ time series with corresponding early warning signals. Early warning signals were calculated using a 20-day rolling window from the onset of winter-like conditions until hibernation began (T_B_ < 33 °C) or the end of monitoring (dotted lines). Hibernation- and Hibernation+ illustrate animals that failed or succeeded in entering hibernation, respectively. (C) Box plots of Kendall’s τ correlation coefficients for each early warning signal. + denotes hamsters that entered hibernation, while - denotes those that did not. (D) 95% confidence intervals for the probability of different types of bifurcation based on the machine learning analysis of pre-hibernation T_B_ dynamics. (E) Within-trajectory discrimination of proximity to hibernation onset at four lead times (20, 15, 10, and 5 days). Upper panels: ROC curves. Lower panels: Box plots of AUC for each model. Each point corresponds to the value for a single hamster. \**Q* < 0.05.

Receiver operating characteristic (ROC) analyses were performed as within-trajectory discrimination analyses at four time-to-onset windows (20, 15, 10, and 5 days) in hamsters with a defined hibernation onset. Changes in T_B_ variance (ΔVariance) produced a higher area under the curve (AUC) than changes in T_B_ deviation from baseline (ΔValue) (**Fig. 3E**). Combining features did not significantly improve performance over ΔVariance alone. Collectively, these results suggest that EWS can track proximity to hypometabolic transitions across species and timescales.

### A mathematical model of mouse thermoregulation exhibits a symmetry-broken pitchfork organization with a saddle-node

We then developed a minimal mathematical model to describe the transition from normothermia to torpor in mice (**Fig. 4A, Methods**). T_B_ is determined by the net balance between heat production (Q_in_) and heat loss (Q_out_), as follows:

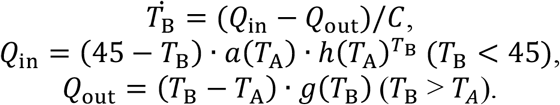

**Figure 4.**
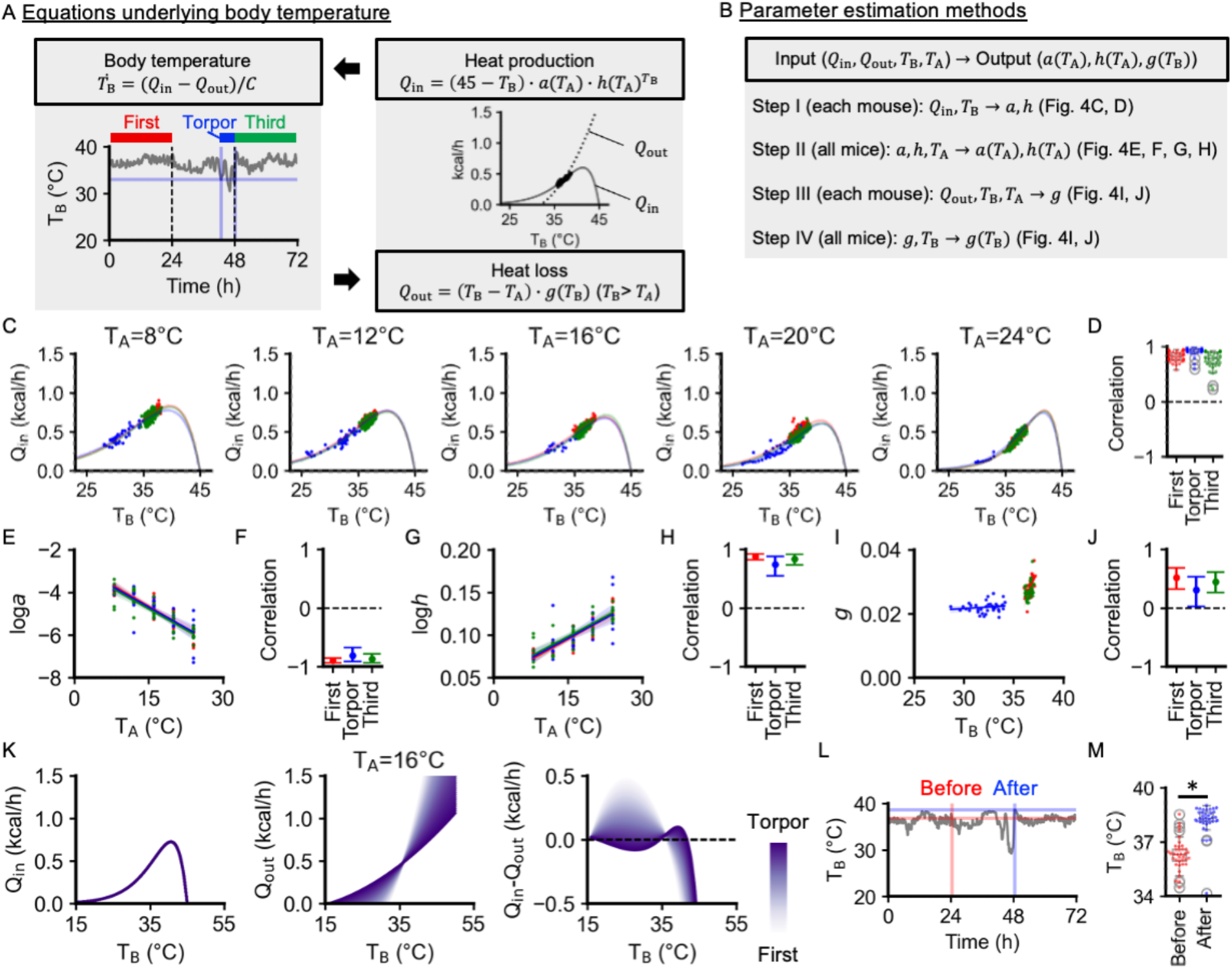
Mathematical modeling of mouse thermoregulation. (A) Model schematic linking heat production (Q_in_), heat loss (Q_out_), core body temperature (T_B_) and ambient temperature (T_A_). Analyses were performed across three periods: First (period of *ad libitum* food access, red), Torpor (period of complete food withdrawal and after T_B_ dropped below the torpor threshold of 33°C, blue), and Third (period of food reintroduction, green). (B) Parameter estimation workflow. For each period, a, h, and g were estimated using Q_in_, Q_out_, T_B_ and T_A_. Q_out_ was not directly measured; interval-mean Q_out_ was inferred from the heat-balance relation, and g was estimated from these inferred values. (C) Representative scatter plots showing the relationship between Q_in_ and T_B_. Curves were fitted separately for the three phases (First, Torpor, and Third) with the equation 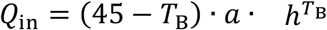. (D) Box plots of correlation coefficients between measured and predicted Q_in_ values. (E) Scatter plots showing the relationship between loga and T_A_, with fitted linear regression lines. (F) 95% confidence intervals for the correlation coefficients between loga and T_A_. (G) Scatter plots showing the relationship between logh and T_A_, with fitted linear regression lines. (H) 95% confidence intervals for the correlation coefficients between logh and T_A_. (I) Scatter plots showing the relationship between g and T_B_, with fitted regression lines. (J) 95% confidence intervals for the correlation coefficients between g and T_B_. (K) Relationship between T_B_ and heat flux components (Q_in_, Q_out_, and Q_in_-Q_out_) at T_A_ = 16°C. Colors are based on the estimated parameter values. (L) An example time course of T_B_, highlighting the maximum T_B_ value during 48–50h (“After,” blue) and the value 24 h earlier (“Before,” red). (M) Box plots comparing T_B_ “Before” and “After”. \**P* < 0.05.

Here, C is the heat capacity; heat fluxes are expressed in kcal h^−1^. Q_in_ peaks below the imposed upper bound of 45°C, at which it falls to zero, whereas Q_out_ is constrained to be non-negative and increases with the temperature gradient T_B_–T_A_. The functions a, h, and g were inferred from measured Q_in_, T_B_ and T_A_, together with an indirect estimate of Q_out_ derived from the heat-balance relation (**Fig. 4B**). First, we fitted individual mouse data to estimate a and h (**Fig. 4B**, step I), achieving significant positive correlations between model predictions and observed values (**Fig. 4C, D**). Second, we quantified the dependence of these parameters on T_A_ (**Fig. 4B**, step II) and found significant log-linear relationships for both a(T_A_) and h(T_A_) (**Fig. 4E-H**). The effective heat-loss coefficient g was then estimated for each mouse (**Fig. 4B**, step III), followed by an analysis of its dependence on T_B_ across mice (**Fig. 4B**, step IV). Although g generally increased with T_B_, the fitted g–T_B_ relationship was shallower in the Torpor epoch than in the First epoch, indicating weaker T_B_ dependence of the effective heat-loss coefficient during torpor (**Fig. 4I, J**).

This analysis resulted in the following mathematical descriptions for First and Torpor states: First:

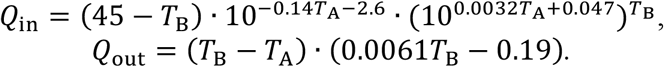

Torpor:

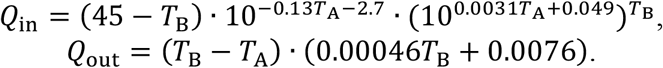

We examined how the phase portrait changed as all fitted parameters were varied simultaneously from their First-state values toward their Torpor-state values (**Fig. 4K, Fig. S1A-D**). This continuation was used to characterize the structural change in the thermoregulatory landscape rather than to reconstruct the temporal trajectory of individual physiological parameters. Because the fitted a(T_A_) and h(T_A_) relations changed only modestly between the First and Torpor endpoint regimes, whereas the g(T_B_) relation changed substantially, the continuation was effectively dominated by the change in effective heat-loss coupling. We therefore interpret the change in g as the principal fitted component associated with landscape reorganization. As a sensitivity analysis, we also held a and h at their First-state values and varied only g toward its Torpor-state relation; this produced the same qualitative phase-portrait reorganization and saddle-node structure. In the First state, the system had a single stable equilibrium around 34-36°C, where heat production and heat loss were balanced. As the parameters approached the Torpor state, the original normothermic equilibrium remained stable and shifted toward higher temperatures. In parallel, a stable low-temperature equilibrium and an intervening unstable equilibrium appeared together. The Torpor parameter set therefore contained three equilibria: a stable torpid state, an unstable state, and a stable high-temperature state. In this landscape, entry into torpor would correspond to movement toward the low-temperature stable state.

We then investigated the relation between this structure and a pitchfork. In an exact pitchfork, a symmetric central state loses stability, and two mirror-image stable states appear. To test whether the fitted model is related to this ideal structure, we expanded the net heat flux near its inflection point (T_c_) into the standard cubic form dy/dt = r(s)y − βy^3^ + ε(s) (**Methods**). The continuation variable s is only an analysis coordinate; it does not represent a measured physiological signal or the actual time course of an animal. The coefficient r crosses zero at s = 0.77 and β is positive, as expected for the symmetric pitchfork parent. However, the symmetry-breaking term ε remains nonzero at that point. Because an exact pitchfork in the cubic approximation requires ε = 0, the fitted model does not pass through an exact pitchfork along this continuation. Instead, symmetry breaking shifts the actual bifurcation to the saddle-node at s = 0.78. The cubic approximation is accurate only close to T_c_, so the saddle-node position was calculated from the full heat-balance model rather than from the cubic approximation (**Methods**). We performed a second analysis to show directly what the asymmetry does. The fitted net heat flux was separated into a symmetric part and an asymmetric part around T_c_. An analysis parameter η controls how much of the asymmetric part is retained: η = 1 exactly reproduces the fitted model, whereas η = 0 removes only the asymmetry without refitting any parameter (**Methods**). The symmetric parent system (η = 0) undergoes a genuine pitchfork bifurcation at s = 0.77, where the central branch loses stability and two symmetric stable branches appear. As the fitted asymmetry is restored, this pitchfork is unfolded and the fitted bifurcation is the saddle-node at s = 0.78. Collectively, the fitted model can be interpreted as an imperfect pitchfork unfolding involving a saddle-node. The pitchfork therefore describes the organizing structure of the landscape, explaining why two stable temperatures become available and why one of them is cold, whereas the saddle-node is the bifurcation encountered along the fitted continuation. Interpreted in this way, the normal-form classifier’s higher probability for the pitchfork class and lower probability for the fold class (**Fig. 2D, 3D**) are not incompatible with the fitted model. This is because the classifier is trained on idealized symmetric forms and is unable to represent an unfolded pitchfork.

To evaluate the model further, we next examined whether several qualitative features of the model were consistent with the experimental data. The model predicted that, during torpor, the upper stable equilibrium would exceed the typical pre-torpor T_B_ (**Fig. 4K**). Qualitatively consistent with this prediction, the maximum T_B_ observed immediately after torpor was significantly higher (38.2 ± 0.8°C, mean ± SD) than the T_B_ observed 24 h earlier (36.3 ± 0.9°C; **Fig. 4L, M**). Of note, because this rebound was observed during recovery after food reintroduction, it should be interpreted as qualitative consistency rather than direct validation of the upper equilibrium in the Torpor parameter regime. At both low and high T_A_, the model showed a single stable equilibrium even in the torpor parameter regime (**Fig. S1E, F**). This qualitative model behavior is consistent with observations that torpor is unlikely to be induced under extreme environmental conditions^7^. Moreover, the model predicted that as T_A_ increases, the slope of the Q_in_-Q_out_ difference curve becomes flatter in the pre-torpor state (**Fig. S1G**), suggesting weaker restoring dynamics and therefore greater variance and autocorrelation in T_B_ and Q_in_. All four measures showed positive associations with T_A_ in the experimental data, three of which reached statistical significance (**Fig. S1H**).

Finally, we estimated the model parameters using only pre-torpor data (**Fig. S1I–R**) to examine how the thermoregulatory landscape changes immediately before torpor onset. As torpor approached, the net heat-flux profile around the normothermic equilibrium became progressively less restoring while the equilibrium remained the only stable state, consistent with the increased variance and autocorrelation observed before torpor onset (**Fig. 2**). Importantly, this change in restoring dynamics was asymmetric away from the equilibrium. At temperatures above the normothermic equilibrium, the magnitude of the net heat flux, |Qin−Qout|, remained relatively large, producing a strong restoring force against upward excursions. In contrast, at temperatures below the equilibrium, |Qin−Qout| progressively decreased, indicating weaker restoration against downward excursions. Thus, even before the low-temperature stable equilibrium emerged, the pre-torpor thermoregulatory landscape became increasingly permissive to finite-amplitude excursions toward lower, but not higher, body temperatures. In the torpor parameter regime, stable equilibria were present at both lower and higher body temperatures. This asymmetric weakening provides a candidate dynamical link to the early-warning signals observed in the body-temperature time series: downward fluctuations encounter progressively weaker restoring forces and can therefore become larger and more persistent while the normothermic equilibrium itself remains stable. Once the saddle-node creates the low-temperature stable state together with an intervening unstable equilibrium, the same directional asymmetry may facilitate excursions toward the newly accessible low-temperature basin. Because stochastic forcing and basin-crossing dynamics were not explicitly modeled, this interpretation is qualitative rather than a quantitative escape mechanism. Collectively, these results suggest a continuous reorganization of thermoregulatory stability as the system approaches the bifurcation, providing a candidate dynamical mechanism that may bias transitions toward the low-temperature torpor state (**Fig. S1S**).

## Discussion

Our results suggest that entry into torpor in mice is associated with a progressive reorganization of the thermoregulatory landscape. In the fitted pre-torpor analyses, restoring dynamics were selectively weaker on the low-temperature side of the normothermic state, coinciding with increased variance and autocorrelation in the body-temperature time series. In the fitted continuation, further reorganization produced a low-temperature stable state through a saddle-node while the normothermic equilibrium remained stable, within an imperfect-pitchfork organization. The fitted pre-torpor landscape became increasingly permissive to downward excursions before the low-temperature state emerged. Once that state becomes accessible, the same asymmetry may bias fluctuations toward the torpor basin. Similar early-warning signatures in Syrian hamsters indicate shared dynamical features across species, although the underlying thermoregulatory mechanism was not directly modeled in hamsters. While gradual declines in T_B_ before hibernation have been documented^3–5,19^, EWS provided greater within-trajectory discrimination of proximity to onset than temperature level alone. Despite its simplicity, our mathematical framework recapitulates several key aspects of mouse thermoregulation, including pre-torpor temperature dynamics (**Fig. 4**), recovery-associated high-temperature rebound (**Fig. 4M**), the T_A_-dependent tendency to enter torpor (**Fig. S1A-F**), and the effects of T_A_ on T_B_ variability during normothermia (**Fig. S1G, H**).

Beyond tracking proximity to the onset of fasting-induced torpor and the first torpor of hibernation, these results suggest a mechanistic hypothesis: the fitted model permits a torpor-like low-temperature state when the effective heat-loss coefficient remains relatively constant as T_B_ falls, while maintaining the coupling between heat production and T_B_. This concept aligns with previous steady-state modeling that inferred minimal changes in the T_B_ set point with reduced heat production^7^, as well as with observations that hypometabolism during daily torpor is primarily due to the effects of low 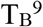. One possible physiological interpretation is that heat-loss pathways, such as vascular tone, become relatively insensitive to changes in T_B_ during torpor; if peripheral vasoconstriction is maintained while heat production increases during arousal, reduced heat dissipation could contribute to the transient elevation of T_B_ observed after torpor.

This possibility remains speculative and requires direct experimental testing. This prediction can guide future research aimed at elucidating the physiological and molecular processes that modulate T_B_ and heat loss during the onset and recovery from torpor. Additionally, EWS may provide a physiological indicator for categorizing the pre-hibernation stage, which could mitigate the confounding factor of individual variability that has complicated molecular studies of pre-hibernation body remodeling. This advantage may be particularly valuable for studying hamster hibernation because there is inter-individual variability in the timing of hibernation onset even under identical housing conditions. To enhance usability, we developed a web application that calculates EWS from user-uploaded body-temperature time series (https://hibernation-tb.streamlit.app/).

In conclusion, EWS track proximity to the onset of torpor and hibernation. In mice, the pre-torpor thermoregulatory landscape becomes progressively permissive to low-temperature excursions, and changes in effective heat-loss coupling are sufficient in the fitted model to extend this reorganization into a bistable landscape through a saddle-node within an imperfect-pitchfork organization. By linking pre-transition fluctuations to directional changes in the thermoregulatory landscape, this work provides a dynamical framework for understanding and experimentally staging transitions into hypometabolism.

## Limitations of the study

The current study has several limitations. First, the biological drivers of the fitted parameter shifts remain unknown. Our mathematical framework is intentionally minimal and phenomenological; it does not incorporate detailed molecular mechanisms, multicompartment heat transfer, or explicit vascular and evaporative control. Second, the First, pre-torpor, and Torpor parameter sets were estimated from separate epochs rather than from a continuously observed physiological control manifold, limiting inference about the path and timing of intermediate parameter changes. Third, technical constraints required using interval-averaged estimates of heat loss rather than direct continuous measurements. Fourth, although hamster trajectories displayed EWS whose classifier-assigned pattern was most similar to a pitchfork class, the absence of heat production records prevented the development of a mathematical model for this species. The underlying mechanisms of the bifurcation may differ between mice and hamsters. Fifth, because the continuation coordinate s linearly interpolates fitted endpoint coefficients and is not a measured physiological trajectory, the numerical saddle-node location depends on the chosen continuation path. In addition, heat loss was not measured directly but inferred from the heat-balance relation, so the inferred change in heat-loss coupling should be interpreted as a model-based effective relation rather than a directly measured causal mechanism.

Sixth, the ROC/AUC analyses quantify within-trajectory discrimination and were not designed to establish out-of-sample predictive performance. Nevertheless, EWS increased as hibernation approached and showed little comparable increase in animals that did not hibernate, supporting evaluation of EWS as a staging marker. Prospective anticipation, robust cutoff values, and predictive accuracy will require validation in larger, independent, and prospective cohorts.

Seventh, the sliding windows used to compute EWS overlap almost completely, so the per-window observations entering the Kendall’s τ trends and the ROC/AUC analyses are not independent. In mice, all animals entered torpor after food withdrawal, and the 24-h pre-torpor windows include data spanning the fed-to-fasted transition. The mouse dataset alone therefore cannot determine whether the EWS specifically encode time-to-torpor or more generally progression after food withdrawal. These are not necessarily alternative explanations, because fasting may itself drive the thermoregulatory system through the observed landscape reorganization. A fasted control group that failed to enter torpor would therefore be required to establish torpor-specificity within mice. Notably, analogous EWS were observed in hamsters without an acute fasting manipulation, and their temporal trends differed between animals that subsequently did and did not hibernate. This cross-species observation argues against the interpretation that the EWS observed here are specific to acute food withdrawal, although it does not establish that the underlying mechanism is identical between species. Eighth, the linear g(T_B_) relations were fitted over a narrow T_B_ range, and the phase portraits extrapolate them well beyond it, so the absolute positions of the equilibria and the numerical value of the saddle-node location are model extrapolations rather than measurements. This limitation primarily affects the global continuation and the positions of remote equilibria; the pre-torpor changes in restoring dynamics around the observed normothermic temperature range are constrained by pre-torpor data and are substantially less dependent on this extrapolation. Ninth, the specific functional forms used for heat production and heat loss are phenomenological assumptions and are not unique. Nevertheless, these functions provided good fits to the measured relationships between heat production, body temperature, and ambient temperature, and the main interpretation of the model concerns the qualitative organization of thermoregulatory stability rather than the exact numerical form of these functions. Combining continuous measurements of heat production and loss with targeted interrogation of vascular, endocrine, and neural pathways in future work should enable richer and more generalizable models of torpor and hibernation.

## Resource availability

### Lead contact

Further information and requests for resources and reagents should be directed to and will be fulfilled by the lead contact, Shinya Kuroda.

## Materials availability

This study did not create any new materials.

## Data and code availability

The data supporting the findings of this study come from the previous studies^3,7^. The code that visualizes the early warning signals is available on our GitHub repository (https://github.com/HikaruSugimoto/BT_app) and the web application (https://hibernation-tb.streamlit.app/).

## Acknowledgments

The images in Figures 2A and 3A are from NIH BioArt (https://bioart.niaid.nih.gov/).

## Author contributions

Conceptualization: HS, SK; Methodology: HS; Investigation: HS; Visualization: HS; Funding acquisition: SK; Project administration: GS, YY, SK; Supervision: SK; Writing – original draft: HS; Writing – review & editing: GS, SN, TS, YY, SK

## Supplementary information

Figure S1

## METHODS

### Key resources table

| REAGENT or RESOURCE | SOURCE | IDENTIFIER |
| --- | --- | --- |
| Deposited data |  |  |
| Previously published mouse fasting-induced torpor data analyzed in this study | Sunagawa and Takahashi<sup>7</sup> | DOI: 10.1038/srep37011 |
| Previously published Syrian hamster hibernation data analyzed in this study | Chayama et al.<sup>3</sup> | DOI: 10.1098/rsos.160002 |
| Experimental models: Organisms/strains |  |  |
| Mouse: C57BL/6J (*Mus musculus*), adult male | Oriental Yeast Co., Ltd. | N/A |
| Syrian hamster (*Mesocricetus auratus*), outbred colony | Japan SLC, Inc. | N/A |
| Software and algorithms |  |  |
| ewstools | Bury<sup>23</sup> | https://github.com/ThomasMBury/ewstools |
| Deep-learning bifurcation classifier | Bury et al.<sup>17</sup> | https://github.com/ThomasMBury/dl_discrete_bifurcation |
| scikit-learn v1.0.2 | scikit-learn developers | https://pypi.org/project/scikit-learn/1.0.2/ |
| SciPy v1.15.2 | Virtanen et al.<sup>24</sup> | https://pypi.org/project/scipy/1.15.2/ |
| A web application that calculates and visualizes early warning signals | This paper | https://github.com/HikaruSugimoto/BT_app;<br>https://hibernation-tb.streamlit.app/ |
| Streamlit | Streamlit | https://streamlit.io/ |
| Other |  |  |
| Temperature-controlled chamber | Shin Factory | Model: HC-100 |
| Thermochron iButton ambient-temperature logger | Embedded Data Systems | Model: DS1922L-F5 |
| Core body-temperature telemetry sensor | DSI | Model: TA11TA-F10 |
| Respiratory gas-analysis mass spectrometer | ARCO System | Model: ARCO-2000 |
| Core body-temperature logger (iButton) | Maxim Integrated | Model: DS1925L-F5 |
| Plasti Dip protective rubber coating for implanted hamster logger | Performix | N/A |

### Animals

This study consists of exploratory analyses of two independent studies that examined fasting-induced torpor in mice^7^ and hibernation in Syrian hamsters^3^. The detailed protocol for the mouse experiments was described previously^7^. Briefly, male C57BL/6J mice were purchased from Oriental Yeast Co., Ltd. and maintained at 21°C and 50% relative humidity on a 12-hour:12-hour light/dark cycle with *ad libitum* access to food and water until experimentation. During experiments, mice were individually housed in temperature-controlled chambers (HC-100, Shin Factory) with continuous ambient temperature monitoring by temperature loggers (Thermochron iButton, DS1922L-F5, Embedded Data Systems). Core body temperature was recorded every 6 minutes using surgically implanted telemetry sensors (TA11TA-F10, DSI) placed in the abdominal cavity under isoflurane anesthesia at least 7 days before the experiments. Metabolic rate was measured continuously by respiratory gas analysis (ARCO-2000 mass spectrometer, ARCO system). All mouse procedures were approved by the Animal Experiment Committee of RIKEN Kobe Institute (approval ID: AH27-05-4) and conducted according to institutional guidelines.

The detailed protocol for the hamster experiments was described previously^3,20,21^. Briefly, hamsters were purchased from an outbred colony (Japan SLC, Inc.) and housed at 24-25°C on a 14-hour:10-hour light/dark cycle with *ad libitum* access to food and water. For core temperature monitoring, temperature loggers (iButton DS1925L-F5, Maxim Integrated) coated with protective rubber (total mass approximately 3.5 g; Plasti Dip, Performix) were surgically implanted in the abdominal cavity under 3-4% isoflurane anesthesia. Data loggers were programmed to record temperature every 10 minutes. After implantation, hamsters were individually housed in polypropylene cages and allowed to recover under summer-like conditions for 1 week before being transferred to winter-like conditions (8-hour:16-hour light/dark cycle, 4°C ambient temperature). All hamster procedures were approved by the Ethics Committees of Hokkaido University (Ethical Approval no. 18-0140) and were conducted according to the ethics guidelines of Hokkaido University.

### Early warning signal assessments

Early warning signals (EWS) were quantified using an established methodology^17,18,22^. Metrics were computed using sliding windows to capture gradual changes in local stability prior to state transitions. Although EWS have been studied in hibernators^8^, several studies have calculated indicators using temperature values extending into the hibernation phase, i.e., after the transition, despite the requirement that EWS be evaluated strictly using pre-transition data. To adhere to this methodological constraint and avoid post-transition contamination, the present analyses restricted EWS computation to pre-torpor/pre-hibernation segments only. For mice, a 24-hour rolling window advanced sample by sample until T_B_ fell below the torpor threshold (33°C). For Syrian hamsters, a 20-day rolling window spanned the interval from the onset of winter-like conditions to either the onset of hibernation (T_B_ < 33°C) or the end of the monitoring period.

Within each window, three indicators were calculated: variance, lag-1 autocorrelation, and the power-spectrum–based index Smax. Trends were summarized using Kendall’s tau (τ). All EWS computations were implemented with ewstools^23^ using its default preprocessing pipeline, which includes detrending, and hyperparameters.

To classify candidate bifurcation types of torpor and hibernation, a previously described deep learning classifier^17^ was applied to body temperature dynamics. Briefly, this model uses a neural network trained on simulated data from different bifurcation models to classify approaching transitions. The neural network architecture combines convolutional neural network and long short-term memory layers (CNN-LSTM). The training data set consisted of 50,000 simulated time series generated by five different models, each representing a different bifurcation type: period-doubling, Neimark-Sacker, fold, transcritical, and pitchfork. Each model combined the normal form of the bifurcation with higher order polynomial terms (up to degree 10) and additive Gaussian white noise. The bifurcation parameter was either linearly increased or held constant (“Null” simulations). The classifiers were trained using Adam optimization with a learning rate of 0.0005 and a batch size of 1024 for 200 epochs. Final predictions were made by averaging the outputs of the classifiers. The classifier was developed for discrete-time bifurcations, whereas the thermoregulatory model used here is continuous-time; accordingly, we used it only as an exploratory pattern classifier. It discriminates among several idealized bifurcation normal forms, one of which is a symmetric pitchfork, and therefore cannot by itself establish that the empirical system has an exact pitchfork symmetry or distinguish a pitchfork from a nearby asymmetric unfolding.

Proximity to onset was evaluated as a within-animal binary discrimination problem at prespecified lead-time horizons. For mice, the horizons were 2.0, 1.5, 1.0, and 0.5 hours before torpor; for hamsters, they were 20, 15, 10, and 5 days before hibernation. For each animal with a defined transition onset, sliding-window observations falling within the specified interval immediately preceding onset were labeled positive, whereas earlier pre-transition observations were labeled negative. For hamsters, ROC/AUC analysis was restricted to animals for which a hibernation-onset time could be defined. Three feature sets were constructed from the same sliding windows: (i) ΔValue, defined as the deviation of T_B_ relative to the first window of the epoch; (ii) ΔVariance, defined as the change in T_B_ variance within the window relative to the first window of the epoch; and (iii) All, comprising both features. Features were z-scored within each animal, and logistic regression was fit separately to each animal and feature set using scikit-learn (v1.0.2). No train/test split was used because the purpose of this analysis was to quantify within-trajectory temporal discrimination rather than out-of-sample generalization. Accordingly, the resulting AUCs were treated as descriptive within-trajectory discrimination measures and not as estimates of prospective predictive performance. For the one-feature models, the logistic transformation is monotonic, so the AUC reflects how consistently the feature ranks time windows close to onset above earlier windows. ROC curves were generated by thresholding the fitted scores over the unit interval to compute sensitivity and 1-specificity, and AUCs were calculated by trapezoidal integration. The same procedure and feature definitions were used for mice and hamsters, with species-specific time horizons.

### A mathematical model of mouse thermoregulation

To characterize the transition between normothermia and torpor states in mice, a mathematical model of mouse thermoregulation was developed. The model was based on the principle that T_B_ is determined by the balance between heat production (Q_in_) and heat loss (Q_out_), as follows:

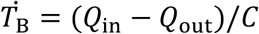

where C is the thermal capacity. Q_in_ and Q_out_ are defined as follows:

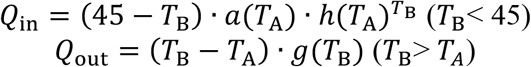

This model included the phenomenological constraint that metabolic heat production declines to zero as T_B_ approaches a fixed upper-temperature cutoff, while heat loss increases with the T_B_−T_A_ gradient. The constant “45” implements this upper-temperature cutoff throughout the analyses; it is a modeling constraint rather than a directly measured denaturation temperature. Substituting nearby values (*e*.*g*., 44 °C) does not alter the qualitative result that the fitted system retains an imperfect-pitchfork organization with a saddle-node; the bifurcation structure and associated early-warning dynamics remain unchanged within physiologically plausible ranges. In contrast, assigning nonphysiological limits (*e*.*g*., values below ~37 °C or extreme values such as 100 °C) violates the model’s biophysical domain and degrades fit and interpretability. For clarity and parsimony, the upper bound was fixed at 45 °C in all reported results. Because a body that is warmer than its surroundings cannot gain heat from them, Q_out_ was additionally constrained to be non-negative over the whole T_B_ range considered. The effective heat-loss coefficient was therefore evaluated as g(T_B_) = max(γ_0_ + γ_1_T_B_, 0), so that Q_out_ = (T_B_ − T_A_)·g(T_B_) ≥ 0 whenever T_B_ > T_A_. This constraint is inactive over the T_B_ range covered by the data, where the fitted g is positive, but it removes non-physical negative heat loss from the extrapolated low-T_B_ portions of the phase portraits.

Importantly, the model does not impose a pitchfork bifurcation or any other specific normal form a priori. Rather, the bifurcation structure emerges from the empirically fitted heat production and heat loss relations. Depending on the fitted coefficients and their continuation, the system may remain with a single stable equilibrium or undergo a fold bifurcation.

This model builds on and extends a previous mathematical model of mouse thermoregulation^7^. The previous model described the relationship between the indices using the following equations:

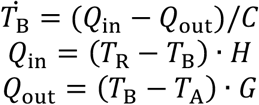

where T_R_ is the set-point body temperature, and H and G are the heat transfer coefficients. Since this model was developed to capture steady-state thermoregulatory mechanisms, the extended model shown above was developed to describe the nonlinear dynamics of mouse body temperature. Moreover, the present model does not impose an explicit set-point temperature T_R_. Instead, T_B_ is treated as fluctuating around temperatures at which heat production and heat loss balance (Q_in_=Q_out_), with stability and variability emerging from the local slope of the net heat-flux function Q_in_−Q_out_. This choice avoids prespecifying a fixed “target” T_B_ and allows the equilibrium (and its stability) to shift dynamically with ambient conditions and parameter changes, thereby representing nonlinear changes in local stability near torpor onset.

Time series were partitioned into three epochs: First (ad libitum feeding), Torpor (food withdrawn and after T_B_ fell below 33°C), and Third (food reintroduced). Parameters *a*(*T*_A_), *h*(*T*_A_), and *g*(*T*_B_) were identified using the following four steps.

Step I: per-mouse estimation of *a*(*T*_A_) and *h*(*T*_A_). Within a given mouse and epoch, T_A_ was constant; consequently, *a*(*T*_A_) and *h*(*T*_A_) could be treated as mouse- and epoch-specific constants. For each mouse and epoch, Q_in_ was estimated from indirect calorimetry, as previously described^7^. Then the relationship

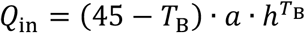

was fit to the paired (T_B_, Qin) observations to obtain *a* and *h*.

Step II: estimation of *a*(*T*_A_) and *h*(*T*_A_). Having obtained *a* and *h* at multiple T_A_ values across mice, the T_A_ dependence was characterized by testing base-10 log–linear models:

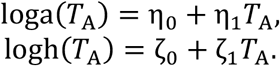

Step III: per-mouse estimation of *g*(*T*_B_). Direct Q_out_ measurements were unavailable. The heat-balance equation 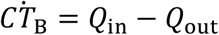 was therefore integrated over intervals [t0, t1] for which the initial and final T_B_ values differed by no more than 0.1 °C; these intervals were treated as approximately returning to the same temperature. Under this approximation, the integrated thermal-storage term was set to zero, giving

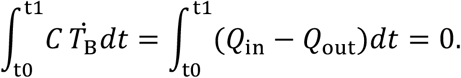

Hence, the time-averaged heat loss within such an interval equals the time-averaged heat production. Parameter estimation used, for each qualified interval [t0, t1], the interval-mean Q_out_ = Q_in_ together with the corresponding interval-mean T_B_ as the pairing for fitting *g*(*T*_B_) in *Q*_out_ = (*T*_B_ − *T*_A_) · *g*(*T*_B_).

Step IV: estimation of *g*(*T*_B_). Because average T_B_ differed across mice, the collection of {T_B_, *g*(*T*_B_)} pairs from Step III was pooled and modeled as a function of T_B_, using linear regression within each epoch:

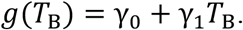

Because the linear g fits are empirical, physiological interpretation was restricted to the data-supported temperature range and to g > 0; extrapolated portions of phase portraits are used only to visualize the mathematical continuation.

More generally, the model is intended as a qualitative, phenomenological description of how the organization of thermoregulatory stability changes between normothermia and torpor, and not as a quantitatively calibrated predictor of body temperature. It is a single-compartment heat-balance formulation with a small number of empirically fitted functions, several of which (in particular the linear g(T_B_) relations, and the a(T_A_) and h(T_A_) relations outside the measured ambient-temperature range) are extrapolated when the phase portraits are drawn. Consequently, only the qualitative structure of the results should be interpreted: the number and stability of the equilibria, the direction in which they move as the parameters are varied, and the existence and ordering of the bifurcations. The specific numerical values reported here (*e*.*g*., the temperatures of the normothermic, unstable and low-temperature equilibria, the temperature at which Q_in_ peaks, the value of the continuation coordinate s at which the saddle-node occurs, and the magnitude of the symmetry-breaking term ε) are model-dependent quantities that are sensitive to the fitted coefficients and to the chosen continuation path. They are reported to make the analysis reproducible and should not be read as physiological predictions of the actual body temperatures adopted by mice, and small numerical differences between these values and the temperatures quoted in the Results text are not meaningful. Importantly, this limitation applies primarily to the global continuation and the positions of remote equilibria. The pre-torpor analysis of restoring dynamics near the normothermic range was performed using parameters estimated from pre-torpor observations and is used to assess changes near the experimentally sampled normothermic regime.

A further limitation is that the temperature dependence of the effective heat-loss coefficient, g(T_B_), was inferred primarily from between-animal variation rather than from repeated measurements spanning a wide range of T_B_ within individual animals. Because each animal contributed an epoch-specific estimate of g, the pooled g– T_B_ relationship does not necessarily imply that g changes with T_B_ in the same manner within an individual animal. We therefore interpret g(T_B_) as an effective, phenomenological relationship rather than as a directly measured within-animal response function. This approach reflects the limited within-animal temperature range over which heat loss could be reliably estimated from the available data. Although this distinction limits a causal interpretation of the fitted g(T_B_) function, the relation remains useful as an empirical effective description for exploring how changes in heat-loss coupling could reorganize thermoregulatory stability. Direct longitudinal measurements of heat loss across a broader range of body temperatures within the same animals will be required to test whether the inferred relationship represents a true within-animal regulatory dependence.

The accuracy of the model fit to the experimental data in the steps I-IV was evaluated using Pearson’s correlation coefficients. The parameters were estimated to minimize residual sum of squares between the model predictions and the experimental data using a nonlinear least squares technique. This analysis was conducted using SciPy (v1.15.2)^24^.

For the phase-portrait analysis between the First and Torpor parameter regimes, a normalized continuation parameter s was used to move all fitted coefficients together from their First-state values (s = 0) to their Torpor-state values (s = 1). The variable s is an analysis coordinate, not a measured physiological control signal or elapsed time. At each value of s, the coefficients defining a(T_A_), h(T_A_), and g(T_B_) were interpolated between their fitted endpoint values.

Equilibrium temperatures were defined by Q_in_ − Q_out_ = 0. An equilibrium was classified as stable when a small temperature displacement produced a net heat flux back toward the equilibrium, corresponding to d(Q_in_ − Q_out_)/dT_B_ < 0, and unstable when the slope was positive. The thermal capacity C changes only the time scale, not the equilibrium positions or this stability criterion.

We quantified local restoring strength as λ = −C^−1^∂(Q_in_ − Q_out_)/∂T_B_; larger positive λ indicates stronger restoration, whereas λ approaching zero indicates weaker restoration. Equilibria were located by bracketing all sign changes of Q_in_ − Q_out_ for T_A_ < T_B_ < 45 °C and then applying bisection. Individual branches were followed in s by Newton iteration from the preceding solution, and saddle-node points were localized by the change in the number of real equilibria.

To determine how the fitted model is related to a pitchfork, we examined the net heat flux F(T_B_, s) = Q_in_(T_B_, s) − Q_out_(T_B_, s) near an inflection point T_c_, where ∂^2^F/∂T_B_^2^ = 0. This choice removes the quadratic term and lets the local dynamics be written in the standard cubic form used to describe a pitchfork. With y = T_B_ − T_c_ and 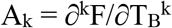 evaluated at T_c_, the Taylor expansion is C dy/dt = A_0_ + A_1_y + (A_2_/2)y^2^ + (A_3_/6)y^3^ + O(y^4^). In the corresponding form dy/dt = r(s)y − β(s)y^3^ + ε(s), r = A_1_/C, β = −A_3_/(6C), and ε = A_0_/C, with A_2_ = 0 at T_c_. An exact pitchfork requires ε = 0. A nonzero ε means that the symmetry is broken and the pitchfork is unfolded. Because the cubic approximation reproduced the full net heat flux to within 2% only for |y| ≤ 2 °C, we used it to characterize the local symmetry breaking, not to locate the bifurcation. Saddle-node points were instead calculated from the full model using F = 0 and ∂F/∂T_B_ = 0. All T_B_ derivatives of F were evaluated analytically. To visualize the effect of the asymmetry without fitting a new model, we separated F around T_c_ into an odd, symmetric component F_odd_(y) = [F(y) − F(-y)]/2 and an even, asymmetric component F_even_(y) = [F(y) + F(-y)]/2. We then defined Fη = F_odd_ + ηF_even_, where η varies from 0 to 1. η = 1 exactly reproduces the fitted model. η = 0 removes only the asymmetric component and gives the ideal symmetric parent system. Thus, η is an analysis parameter used to reveal the symmetry-breaking structure; it is not a physiological quantity.

Importantly, the saddle-node in the fitted continuation creates a low-temperature stable state while the normothermic equilibrium remains stable; the deterministic model therefore does not, by itself, specify what triggers movement from the normothermic basin to the torpid basin. The observed fluctuations in T_B_, and potentially unmeasured fluctuations in physiological variables, could contribute to crossing the intervening basin boundary once the low-temperature state becomes accessible. The sources, magnitude, and physiological triggers of such fluctuations remain unresolved and require future investigation. Accordingly, we interpret this as a qualitative stochastic transition mechanism rather than a quantitative escape model.

## Server implementation

A web application that calculates and visualizes early warning signals (https://hibernation-tb.streamlit.app/) was developed using Streamlit (https://www.streamlit.io). The code is available on GitHub (https://github.com/HikaruSugimoto/BT_app).

## Statistical analysis

Matched within-animal comparisons, including comparisons of AUCs obtained from different feature sets and the Before–After comparison of body temperature, were performed using paired t-tests. Welch’s t-tests were used for comparisons between independent groups. Per-animal Kendall τ values were tested against zero using two-sided one-sample t-tests. When multiple related comparisons were performed, P values were adjusted using the Benjamini–Hochberg procedure^25^. Ninety-five percent confidence intervals for classifier probabilities, group ROC curves, and across-animal correlation summaries were estimated by nonparametric bootstrap resampling of animals with 10,000 replicates. Associations between variables were assessed using Pearson’s correlation coefficients, and temporal trends in EWS were summarized using Kendall’s τ. Statistical significance was set at *P* < 0.05 or *Q* < 0.05. Statistical testing was conducted using SciPy (v1.15.2).

---

**Figure S1.**
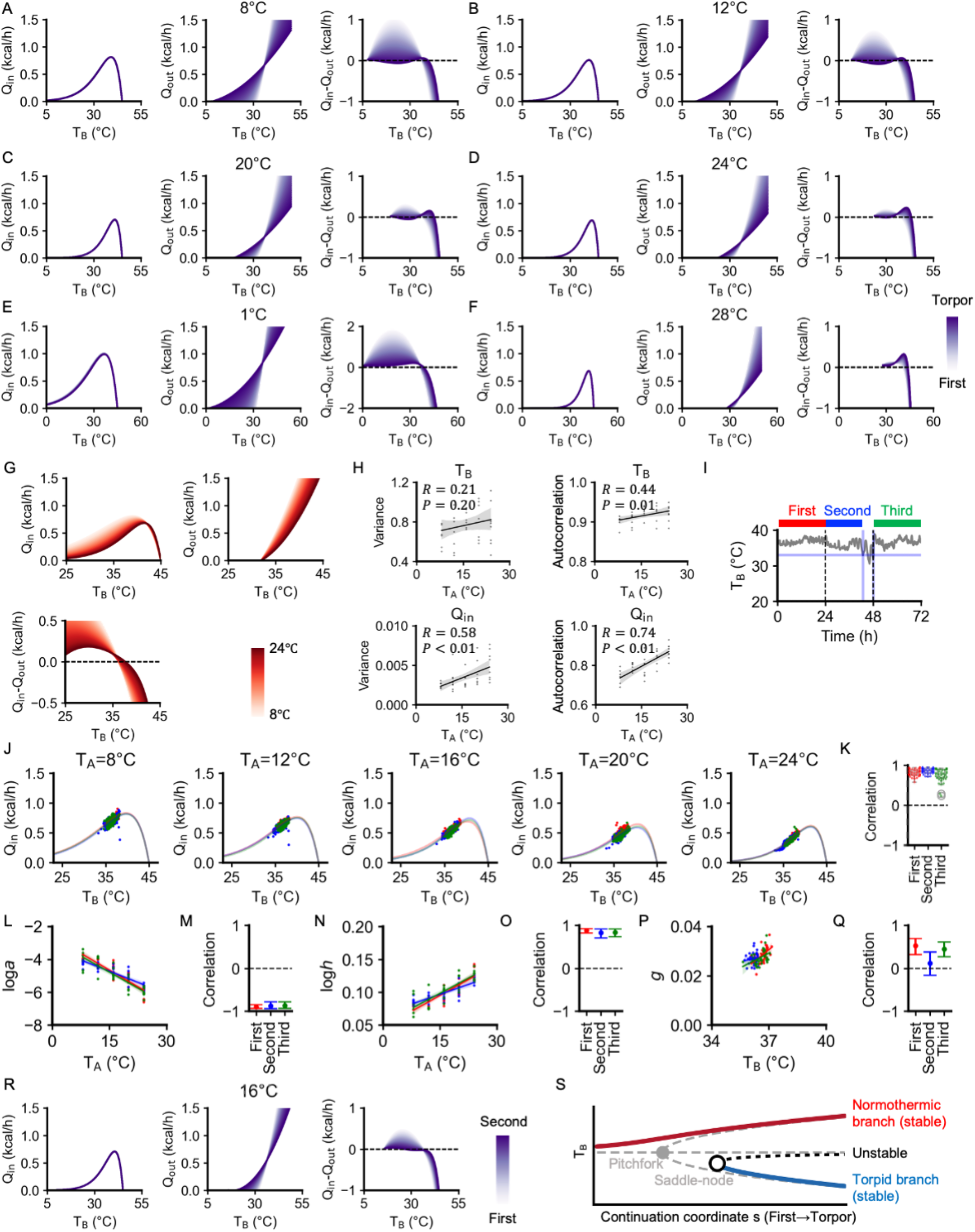
Mathematical analyses of mouse thermoregulation. (A-F) Relationship between T_B_ and heat flux components (Q_in_, Q_out_, and Q_in_-Q_out_) at six different T_A_: 8°C (A), 12°C (B), 20°C (C), 24°C (D), 1°C (E), 28°C (F). Colors denote parameter values. (G) Relationship between T_B_ and heat flux components (Q_in_, Q_out_, and Q_in_-Q_out_) during the first 24-hour period. Colors are based on T_A_. (H) Scatter plots with fitted linear regression lines for T_A_ versus four thermoregulatory parameters (T_B_ variance, T_B_ autocorrelation, Q_in_ variance, and Q_in_ autocorrelation). Each point corresponds to the values for a single mouse. Gray shaded areas indicate the 95% confidence intervals. (I) Representative time course of T_B_. The following analysis includes three periods: initial ad libitum feeding period (first, red), food withdrawal period before torpor onset (T_B_ > 33°C; second, blue), and food reintroduction period (third, green). (J) Representative scatter plots showing the relationship between Q_in_ and T_B_ in individual mice, with curves fitted using the equation 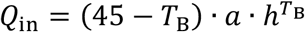. (K) Box plots of correlation coefficients between measured and predicted Q_in_ values. (L) Scatter plots showing the relationship between loga and T_A_, with fitted linear regression lines. (M) 95% confidence intervals for the correlation coefficients between loga and T_A_. (N) Scatter plots showing the relationship between logh and T_A_, with fitted linear regression lines. (O) 95% confidence intervals for the correlation coefficients between logh and T_A_. (P) Scatter plots showing the relationship between g and T_B_, with fitted regression lines. (Q) 95% confidence intervals for the correlation coefficients between g and T_B_. (R) Relationship between T_B_ and heat flux components (Q_in_, Q_out_, and Q_in_-Q_out_) at T_A_ = 16°C. Colors are based on the estimated parameter values. (S) Bifurcation diagram of the fitted model, showing how the equilibrium T_B_ depends on the model parameters at T_A_ = 16 °C. Equilibria are shown as the parameters are moved from the First set (s = 0) to the Torpor set (s = 1) along the normalized continuation coordinate s; s is an analysis coordinate, not elapsed time and not a measured physiological signal. Solid lines are stable equilibria (red, normothermic branch; blue, torpid branch) and the black dotted line is the intervening unstable equilibrium. For s < 0.78 the system has a single stable equilibrium. At s = 0.78 a stable and an unstable equilibrium are created together at a saddle-node (open circle), so a torpid state becomes available while the normothermic state remains stable and moves to higher temperatures. Grey dashed lines show the corresponding symmetric parent system, obtained by removing the asymmetric (even) component of the net heat flux about the expansion point Tc as described in the Methods; that system has an exact pitchfork at s = 0.77 (grey circle), at which the central branch loses stability and two branches appear. The offset between the two markers is the effect of the symmetry-breaking term: the pitchfork organizes the landscape and accounts for why two stable temperatures become available and why one of them is cold, whereas the saddle-node is the bifurcation present in the fitted system. The grey dashed branches are drawn only over the range in which the cubic expansion about Tc is reliable. As stated in the Methods, the model is qualitative: the number, stability and ordering of the equilibria, rather than their exact temperatures, are what should be interpreted.

